# Estimating information flow through a memory system: the utility of meta-analytic methods for genetics

**DOI:** 10.1101/020586

**Authors:** Tugce Yildizoglu, Jan-Marek Weislogel, Farhan Mohammad, Edwin S.-Y. Chan, Pryseley N. Assam, Adam Claridge-Chang

## Abstract

Critics of significance testing claim that this statistical framework promotes discrepancies by using arbitrary thresholds (α) to impose reject/accept dichotomies on continuous data, which is not reflective of the biological reality of quantitative phenotypes. Here we explore this idea and evaluate an alternative approach, demonstrating the potential for meta-analysis and related estimation methods to resolve discordance generated by the use of traditional significance tests. We selected a set of behavioral studies proposing differing models of the physiological basis of *Drosophila* olfactory memory and used systematic review and meta-analysis approaches to define the true role of lobular specialization within the brain. The mainstream view is that each of the three lobes of the *Drosophila* mushroom body play specialized roles in short-term aversive olfactory memory [1-5], but a number of studies have made divergent conclusions based on their discordant experimental findings [6-8]. Multivariate meta-regression models revealed that short-term memory lobular specialization is not in fact supported by the data, and identified the cellular extent of a transgenic driver as the major predictor of its effect on short-term memory. Our findings demonstrate that meta-analysis, meta-regression, hierarchical models and estimation methods in general can be successfully harnessed to identify knowledge gaps, synthesize divergent results, accommodate heterogeneous experimental design and quantify genetic mechanisms.

## LAY SUMMARY

Significance testing is a statistical method widely used to assess the effect of gene variation, particularly in genetic models e.g., mice and flies. While many traits vary continuously, significance testing is designed to produce a simple yes/no outcome that is unsuitable for smoothly varying effects. For decades, statistical texts have proposed that significance testing has a distorting effect. We examined the influence of significance testing in vinegar fly research on short-term olfactory memory, an extensively-studied quantitative phenotype in a major model genetic system. Significance tests have been previously used to show that a particular brain sub-region - the gamma lobe - is highly specialized for short-term memory. We re-analyzed published data using estimation methods that place emphasis on quantification: meta-analysis and hierarchical models. We show that the gamma lobe shares memory processing roles with two other brain lobes. Using neuronal cell count data, we also show an absence of even relative specialization, revealing that memory processing is distributed across neurons in all three lobes. We propose that significance testing distorts other areas of genetic analysis in similar ways; our data indicate that this can be ameliorated with the adoption of estimation statistics instead.

## INTRODUCTION

Contradictory results from research are commonplace. Discordance stems from sampling error and methodological differences, both sources of variability that are largely unavoidable. One concern is the widespread acceptance of weak significance testing power; a recent report revealed that many neuroscience studies have a test power below 40% [9], and the same pattern is likely to be seen across other disciplines. However, critics of significance testing itself claim that this statistical framework needlessly accentuates differences. The numerous conceptual and practical limitations of significance tests [10] include the inherent volatility of p-values, even when there is moderate statistical power [11]. Simulation demonstrates that test results from different studies can easily be discordant due to sampling error alone, even assuming flawless methodological standardization [11]. Moreover, significance testing may exacerbate discordance by using an arbitrary threshold to force a binary outcome (reject/accept) from continuous data [12]. To illustrate, a pair of alpha 0.05 tests on two replicated experiments with identical effect sizes could produce p-values of 0.049 and 0.051: in the significance testing framework these results are starkly discordant, when in reality the biological outcome is all-but the same [12]. The arbitrary reject/accept dichotomy might also lead to the false impression that a substantial (but non-statistically significant) effect is irrelevant. Conversely, a highly powered sample size could give the misleading impression that a minuscule (but statistically significant) effect is of great importance [11]. Thus, some consider that fields relying solely on significance testing to draw their conclusions are particularly susceptible to discrepancies and may be incapable of resolving apparent irreproducibility.

In medical research, the complementary methods of systematic review and meta-analysis are routinely used to synthesize evidence from multiple studies and to reconcile divergent findings [13]. However, such approaches are rarely applied to basic research fields like neuroscience. Taking a mainstream sub-field of neuroscience as an example, a PubMed search in late 2014 with the phrase “meta-analysis AND (learning OR memory) AND mouse” identified fewer than ten studies in a field of >35,000 articles. We therefore decided to ask whether meta-analytic methods could be used to evaluate the possible influence of significance testing dichotomization. In seeking a suitable research field we required an unresolved hypothesis for which the published studies included adequate sample sizes for meaningful analysis, and used a standardized protocol so that the data would not be dominated by sampling error (weak statistical power) and methodological heterogeneity. We selected the investigation of the neuronal mechanisms of olfactory memory in *Drosophila melanogaster*. Olfactory memory in *Drosophila* is measured using the classical T-maze olfactory conditioning assay, where groups of flies are conditioned by pairing an odor with an electric shock and subsequently assessed for their ability to avoid the conditioned odor when given a choice of two different odors presented at the end of the maze arms. A particular strength of the T-maze is its use of hundreds or thousands of animals in a single experiment [14], which helps to minimize the sampling error that is often inherent in rodent assays and assays using smaller numbers of insects [15,16]. In addition, both the T-maze apparatus and the training regime is largely standardized between labs [14].

Thirty years of T-maze experiments have elucidated many of the genetic, molecular and neural mechanisms of olfactory learning [1-5,17]. A landmark study showed that restoring the adenylyl cyclase gene *rutabaga (rut*) to a brain structure called the mushroom body is sufficient for short-term olfactory memory [6], connecting memory formation to cyclic adenosine monophosphate-mediated plasticity [18]. Experiments using inhibition of synaptic transmission by temperature-sensitive *shibire (shi)* [19-21] showed that neurotransmission from the mushroom body is essential [20,22]. Targeted expression of genes in specific neuronal circuits is possible with the use of transgenic ‘driver’ lines [23]. Manipulations based on *rut* restoration and *shi* inactivation form the foundation of a large number of studies aiming to further define the role of the mushroom body in olfactory learning. The mushroom body itself exists as three anatomically distinct lobes, αβ, α’β’, and γ [24]; studies on middle- and long-term memory (MTM and LTM) have revealed distinct lobe requirements in the different memory phases [21,25-28]. However, the three lobes’ specializations remain unclear when it comes to short-term memory (STM). While the mainstream view is that *rut* activity in the γ lobes is sufficient to rescue STM [8], some studies have alternately concluded that *rut* restoration can only partially rescue [7], or is merely of importance to STM [6]. There is similar controversy on the role of *rut* activity in the αβ lobes, with *rut* restoration said to have either no effect [8], or to partially rescue STM for certain odors [7].

In the present study, we aimed to evaluate the mainstream view that there is strong lobular specialization of STM function in the mushroom body, and to assess the extent to which the varying perspectives on this subject resulted from significance testing’s forced dichotomization. Using meta-analytic methods, we examined the proposals that restoration of *rut f*unction to the γ lobes alone is sufficient to rescue wild type STM and that only *shi* function in the γ lobes is necessary for STM. In both cases, meta-analysis of published studies spanning more than a decade found no evidence for strong lobular specialization. A subsequent analysis with multi-level meta-regression, an advanced estimation technique, revealed that numbers of mushroom body cells explained nearly all transgenic effects. These results confirm claims made by statistical texts that systematic review, meta-analysis and related estimation methods can be applied to resolve currently conflicting data and give new quantitative perspectives. In addition to its role in review, we conclude that routine use of both basic and advanced estimation methods would aid the planning, analysis and interpretation of research.

## RESULTS

### Systematic literature review of *rutabaga* and *shibire* interventions in short-term aversive olfactory memory

The review yielded ten studies that fulfilled the criteria (Figure 1A). Seven studies contained 81 experiments related to *rutabaga* restoration **[6-8,22,29-31]**, with a total of 748 experimental iterations and 745 control iterations (see Table 1). Each iteration is the mean of two half-PI scores, which typically each use 50-100 flies, thus representing an estimated total of 150,000-300,000 assayed flies. Table 1 also lists the 5 studies that contained 37 experiments related to *shibire*-mediated inactivation **[7,20,21,25,29]**, 263 experimental iterations and 265 control iterations, giving a total of 50,000-100,000 flies.

**Figure 1.**
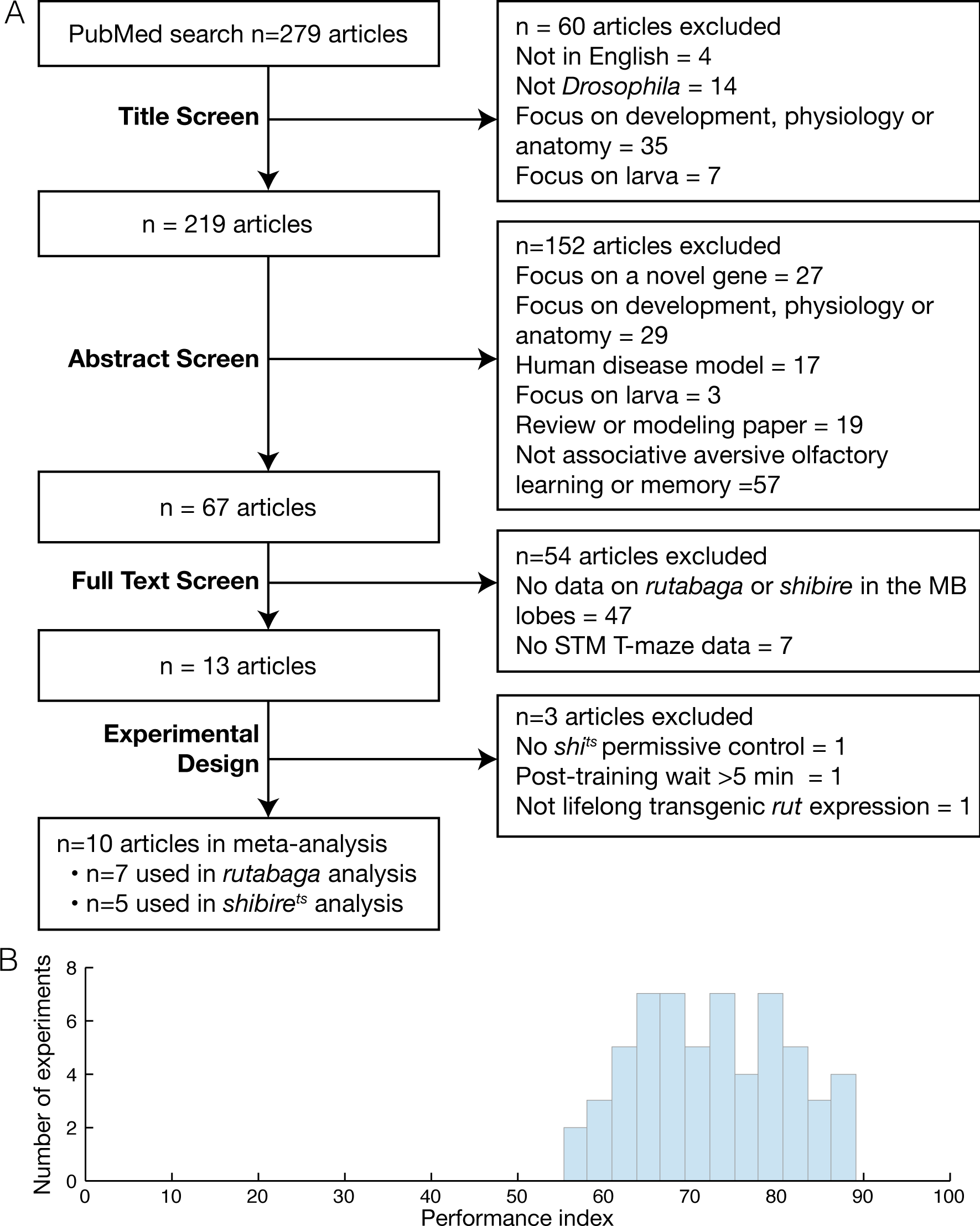
Review overview. **A.** Flow chart of systematic literature review procedure. The literature was reviewed in a five stage process, starting with a PubMed search that yielded 279 articles, followed by four screens of increasing detail, reviewing the article title, abstract full text and experimental design. A total of ten articles, two of which included relevant data for both rutabaga and shibire^ts^ experiments, were used in the meta-analyses. **B.** Histogram of performance indices for all control experiments identified by the review.

**Table 1.**
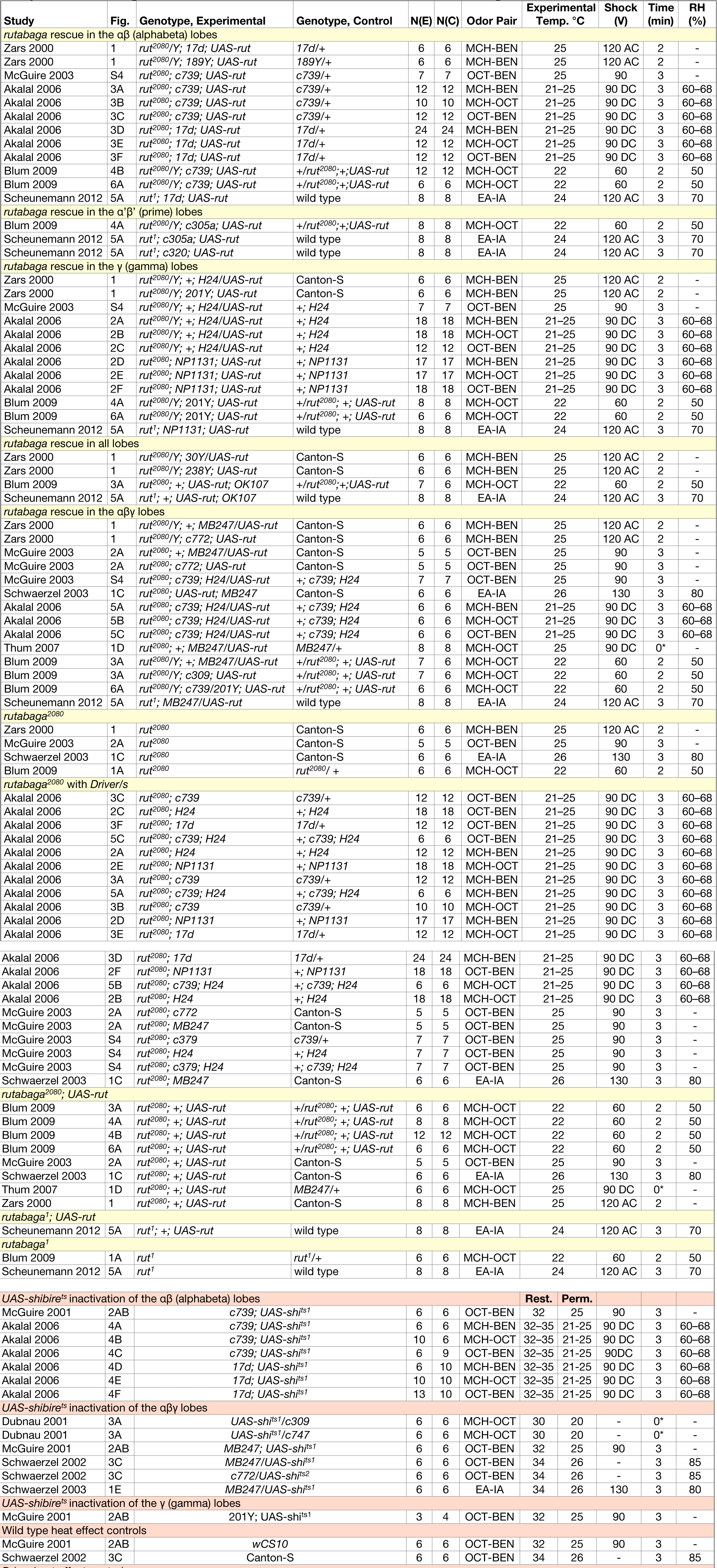
Characteristics of included experiments. All experiments are listed and identified by their study, figure panel and genotype/s. We name the most precise genotype possible based on the information given in the original article. Odor pair, range experimental temperature or temperature range, the nature of the conditioning shock and the relative humidity (RH) are also listed. The time delay between training and testing is listed in minutes; those labelled ‘0*’ were reported as following training ‘immediately.’ Shock is listed in volts; current type is omitted if not reported in the original study. Cells containing a dash indicate that the information was not found in the original article.

### Experimental variability

Despite standardization of aspects of the T-maze, some methodological variation between studies was observed, including different control genotypes, varying odor pairs, temperatures, shock voltages, humidity and post-training delay times prior to testing (Table 1). These differences, along with other uncontrolled variables common to behavioral experiments, would explain the variability seen in data from control experiments (Figure 1B). We found considerable heterogeneity in several of the meta-analyses. In the six *rut* analyses, overall heterogeneity was low in three (I^2^ < 50%), and high in three (I^2^ > 75%); subgroup heterogeneity (i.e. variance due to genotype differences) was low in four, and high in two (Figures 6-11). In the *shi* analyses, overall heterogeneity was high in two and moderate in one, while their subgroup heterogeneity values were 34%, 64% and 80%.

### *Rutabaga* function is required for 60% of wild type learning

We aimed to estimate the learning contribution made by restoring *rutabaga* function to each of the three lobes. The meta-analyses on *rutabaga* experiments produced 6 meta-analytical estimates of the effects of manipulating *rut* in the mushroom body lobes (Figure 2B). Data pooled from *rut*^*1*^ and *rut*^*2080*^ reveal that the strong *rut* hypomorphic alleles reduce learning to 40% of wild type (-60% [95CI -56, -64]) (Figure 2B, 6). The forest plot in Figure 2A illustrates the individual effect sizes from 36 experiments and pooled effect sizes of the *rut* alleles (complete forest plot is shown in Figure 6). The data exhibit substantial overall heterogeneity (I^2^= 76%) and genotype subgroup heterogeneity (I^2^=88%). This heterogeneity may derive from the methodological variation noted above, but in the case of the strong *rut* alleles we note that the weakest effect is seen in the *rut*^*2080*^; *UAS-rut* subgroup (-45% [95CI -38, -52]), suggesting leaky expression from the transgene as one possible source (i.e. expression from the *UAS-rut* transgene independent of GAL4 transcriptional activation).

**Figure 2.**
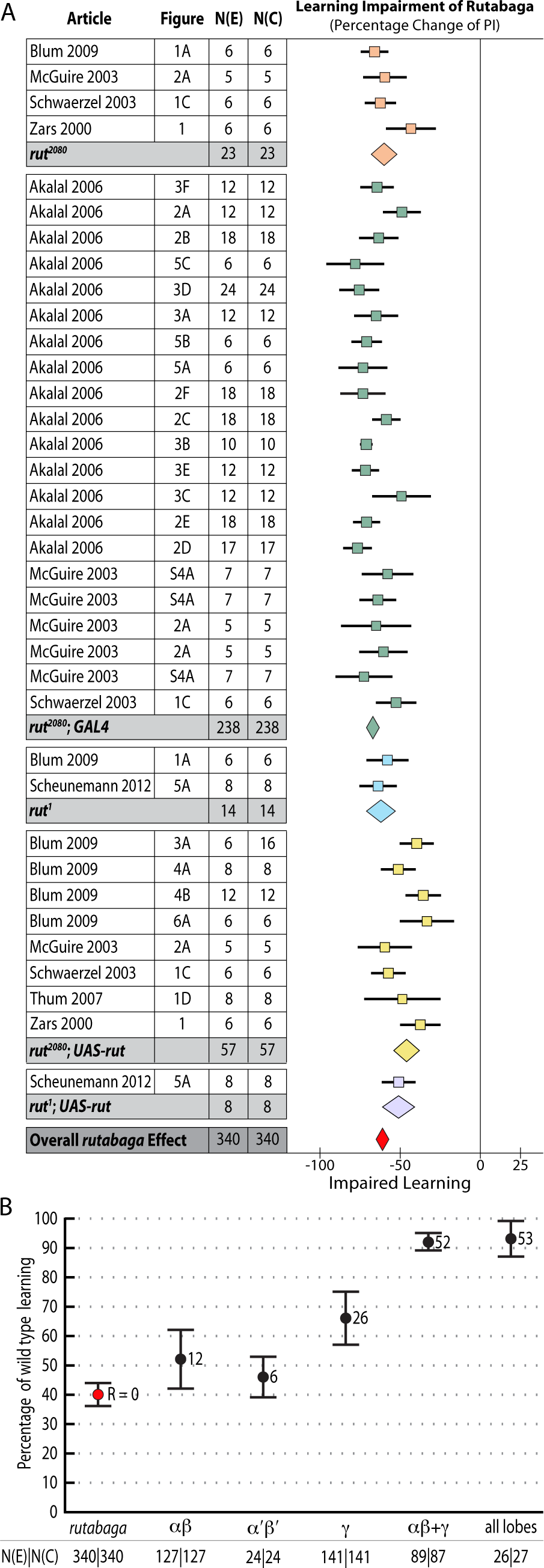
Meta-analyses of *rutabaga* mutant lines and targeted transgenic restoration. Short-term memory data are expressed as percentages. **A.** A summary forest plot of learning changes observed in 340 experiments with *rut* mutant lines, with subgroups showing the differences between the various *rut* alleles and strains. Learning is expressed as a percentage change relative to wild type. The red diamond on the bottom line indicates that the overall impairment in learning in the *rut* hypomorphs relative to wild type controls is -60% [95CI -56%, -64%]. The complete forest plot is given in Figure 5. **B.** Summary estimates from the *rut* mutant meta-analysis and five meta-analyses of lobular restoration experiments. Learning is displayed as a percentage of wild type learning. The markers indicate the proportion of learning relative to wild type expressed as a percentage; error bars are 95% confidence intervals. To the right of the markers are numbers for the amount of rescue (R =) relative to the rut hypomorphs. N(E) and N(C) are the experimental and control iterations respectively. Except for the α’β’ lobes (p=0.17), all lobe categories showed a statistically significant partial rescue of learning (αβ p=0.029, γ p<1 × 10-45, αβ+γ p=1.1 × 10-16, all lobes p<1 × 10-45) when compared with rut learning.

### *Rutabaga* restoration to the γ lobes rescues 26% of wild type STM

Some studies have reported that complete rescue requires *rut* restoration in both αβ and γ lobes **[7]**, while others report that restoring *rut* activity in the γ lobe is sufficient to rescue STM, and that the αβ lobes’ *rut* activity has little or no STM role **[8]**. We used the meta-analytic data to specifically examine the lobular specialization hypothesis (Figures 5-10). The overall *rutabaga* loss-of-function effect was used as a reference point to which we compared the lobe restorations, shown in Figure 2B. Restoring *rut* function to each of the lobes revealed partial rescue: α’β’ rescues by 6% [95CI -1.5, 13.5], αβ rescues by 12% [95CI 2, 22] and γ rescues by 26% [95CI 17, 35]. When *rutabaga* was restored to both the αβ and the γ lobes, memory was rescued by 52% [95CI 50, 55]. Restoring *rutabaga* to all three lobes gave only 1% additional improvement (53% [95CI 47, 59]) compared to the rescue in the αβ + γ lobes, therefore *rut* in the α’β’ cells appears to have a minor effect on STM. Of the enhancer trap drivers included in the γ meta-analysis, 201Y contains a minority of αβ cells **[32]**. A variant analysis that removed 201Y from the γ group and reassigned it to the αβ + γ group resulted in weaker effects for both: only 20% [95CI 10, 31] γ rescue, while αβ + γ rescue was reduced to 49% [95CI 46, 52]. Taken together, these results are incompatible with the hypothesis that restoring *rut* activity to the γ lobe alone is sufficient to rescue the *rut*^*-*^ phenotype. From the lobe perspective, we conclude that normal STM requires *rut* function in both αβ + γ lobes.

### Heating flies above 30°C impairs short-term memory

Using the temperature-sensitive alleles of *shibire* to block neurotransmission requires heating flies to over 30°C, which can lead to additional heat-related effects **[28]**. Researchers accommodate this possibility with separate ‘heat control’ flies that do not express *shi*^*ts*^. We estimated the magnitude of this effect by meta-analysis, shown in Figure 3A (complete forest plot in Figure 11). Data pooled from 23 such experiments with three types of genotype (wild type, *Driver-GAL4/+* and *UAS-shi^ts^/+*) revealed that the overall effect of heating flies from the permissive temperature (20-26°C) to 30-35°C is a 17% [95CI 12, 22] reduction in memory. This decrement can be expected to affect the *UAS-shi^ts^* inactivation data from the same studies, so we used 83% of wild type memory in Figure 3B as the zero reference point to estimate the specific effects of lobe inactivation.

**Figure 3.**
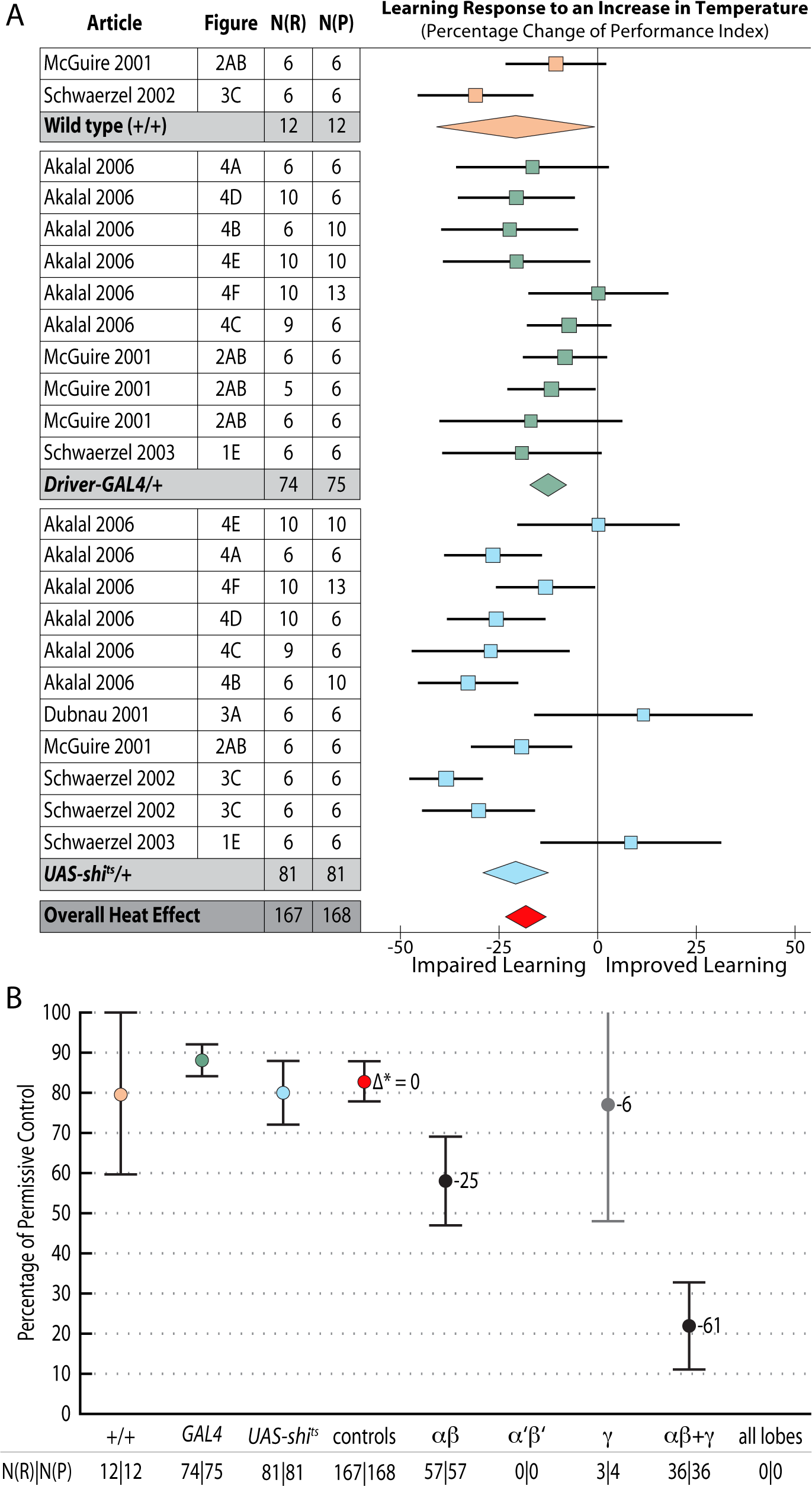
Meta-analyses of *shibire*^*ts*^ inhibition of neurotransmission in the mushroom body lobes. Learning data are expressed as percentages. **A.** A summary forest plot of learning changes in heat treatment controls, with subgroups showing the differences between 3 types of controls. Learning is expressed as a percentage change relative to wild type. The red diamond on the bottom line indicates that the overall impairment in learning in flies exposed to elevated temperature is -17% [95CI -12%, -22%]. A complete forest plot is shown in Figure 11. **B.** Summary estimates from the heat exposure controls and three meta-analyses of lobular inactivation experiments. Colored markers correspond to diamonds in panel A. Learning at the restrictive temperature is shown as a percentage of learning at the permissive temperature; error bars are 95% confidence intervals. To the right of the markers are numbers learning impairment (Δ* =) relative to the synthetic heat effect control. N(R) and N(P) are the restrictive and permissive iterations respectively. The αβ lobes (p=0.0001) and the αβ+γ combination (p<1 × 10-45) show statistically significant impairment while the γ lobes do not (p=0.7071). The γ lobe bar is in grey as it derives from only a single experiment with few replicates. There were no data in the literature on the α’β’ lobes or drivers that encompass all mushroom body lobes.

### Neurotransmission from the αβ + γ lobes accounts for 61% of STM

Drivers that express in both the αβ and γ lobes reduced performance by 61% [95CI 50, 72] relative to heated control flies. Inactivating the αβ lobes produced a 25% [95CI 14, 37] reduction in STM. The best estimate for γ lobe inactivation is a 6% reduction [95CI 35% reduction, 24% increase] relative to heated controls. This γ lobe estimate appears to be negligible, but has very wide confidence intervals and is drawn from only a single experiment with three iterations. Surprisingly, the literature review found no <5 min STM data on the impact of *shibire*^*ts*^ inactivation of either the entire mushroom body (All lobes) or the α’β’ lobes (empty columns in Figure 3B); at the time of the review the only studies reporting results for these interventions examined later memory, at 15 min or beyond **[28]**. The substantial decrement in the αβ lobe inactivation experiments (25% reduction) is incompatible with the idea that this lobe plays only a negligible role in STM. The paucity of data for γ, α’β’ and All lobes in STM highlights an area that would benefit from future experimental attention.

### Cell number accounts for the majority of driver variation

Observing high heterogeneity (I^2^) in some of the meta-analyses, we attempted to identify the source of variability, and examine the original hypothesis from a different perspective. Electrophysiological evidence **[33]** and anatomical connectivity analysis **[34]** indicate that the Kenyon cells, the intrinsic neurons of the mushroom body, are randomly connected to their olfactory input neurons. The lack of structured connectivity suggests that, for some or all odor-related functions, individual Kenyon cells are interchangeable; thus raising the possibility that a cell’s lobular identity might be less important than its participation in a stochastically nominated odor-responsive ensemble. As three of the seven relevant meta-analyses showed driver heterogeneity as accounting for more than half of their variance, we asked whether the number of cells captured by a driver could explain some of the unaccounted variance. We extracted cell count data from an anatomical study that counted Kenyon cells for many of the drivers **[32]**. The driver-specific meta-analytic STM estimates were subjected to an initial simple linear regression against the drivers’ available cell counts in both *rut* restoration and *shi*^*ts*^ inactivation. These indicated that cell numbers accounted for about 80% of the driver memory variance (*rut* R^2^ = 0.79 [95CI 0.39, 0.94], p=2.5 × 10^−4^; *shi*^*ts*^ R^2^ = 0.77 [95CI 0.14, 0.96], p=8.4 × 10^−3^). As simple linear regression is unable to account for the full complexity of such hierarchical data, we constructed hierarchical, multivariate, weighted meta-regression models accommodating other variables that might explain some of the variance induced by differences in experimental design. These models were also able to account for the clustering of experiments within studies and for the shared control design in *rut* experiments, and included weighted estimates for each driver by the number of contributing experiments (described fully in Methods). The hierarchical meta-regression model of *rut* showed a strong relationship with driver cell count, generalized-R^2^ = 0.84 [95CI 0.79, 0.89] (Figure 4A). The meta-regression model of *shi* data similarly revealed a large effect size for the cell count relationship, generalized-R^2^ = 0.88 [95CI 0.84, 0.92] (Figure 4B). Compared with simple linear regression, the hierarchical models revealed stronger trends with substantially improved precision. These results are incompatible with the strong lobular specialization hypothesis of *rut* and *shi* function. Rather, drawing on data from thousands of T-maze iterations (N = 1008, 1006) while accounting for experimental heterogeneity, they constitute compelling evidence that each driver’s extent of neuronal expression can account for the majority of that driver’s short-term memory effect.

**Figure 4.**
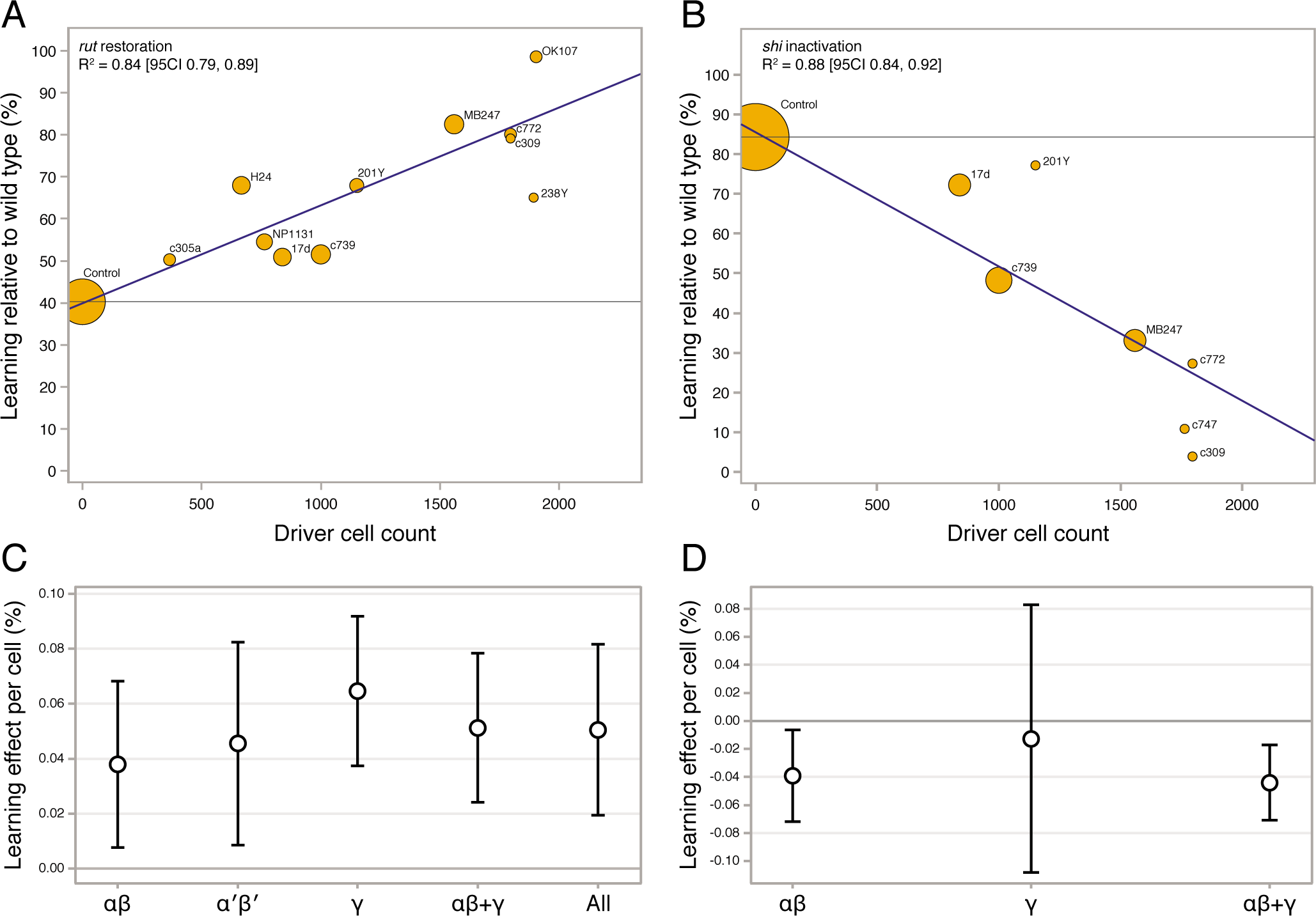
The extent of drivers’ Kenyon cell expression accounts for the majority of short-term olfactory memory effects. The estimated Kenyon cell counts for drivers were taken from Aso et al. 2009. The memory effect sizes are derived from nested, weighted, multivariate meta-regression models that adjusted for confounding variables that contributed to heterogeneity. **A.** Bubble plot of *rut* restoration; the cell count of driver lines accounts for 84% of the variance of the learning effects of rut restoration (p < 0.0001). Each bubble’s area indicates that estimate’s weight in the regression model; the blue fit line has a slope of 0.023% per cell [95CI 0.016, 0.030]. The grey line indicates the level of no rescue, i.e. the learning level of *rut* mutants. **B.** For *shi*^*ts*^ inactivation, 88% of the learning variance is attributable to the number of cells encompassed by the driver (p < 0.0001). The blue fit line has a slope of -0.034 % per cell [95CI -0.046, -0.0216]; the grey line indicates the level of no effect, i.e. the learning expected from the effect of heat alone. **C.** Learning effect per cell in mushroom body sub-regions from *rut* restoration in different lobes and combinations, adjusted for heterogeneity effects. Error bars are confidence intervals; there are no statistical differences between rut lobe categories. **D.** The *shi*^*ts*^ learning effect per cell in two lobes and their combination. There are no statistical differences between *shi*^*ts*^ lobe categories.

### Kenyon cells in different lobes make equivalent contributions to STM

Different Kenyon cell drivers’ varying impact on learning is primarily a result of how many cells they are expressed in: cell count as the overwhelmingly dominant factor therefore excludes highly specialized roles for *rut* and *shi* in different lobes’ Kenyon cells. However, it is possible that minor quantitative differences explain the remaining unaccounted for 12-16 % of STM variance in the meta-regression models. Within the overall memory-cell count trend in Figure 4A, several drivers’ estimates do not fall on the regression line. To account for such deviations from the overall cell number trend, we aimed to factor out cell number and focus specifically on the potency of each neuron captured by a driver. We built new models in which the learning effect size of each driver line was first divided by the number of expressing cells, and weighted hierarchical meta-regression models were then used to perform synthesis by lobular category. These models produced estimates of a typical Kenyon cell’s effectiveness within each lobe category (Figure 4C & D). The *rut* rescue-per-cell data and the *shi* loss-per-cell data both show that there are no substantial differences between any lobe categories. In summary, when cell numbers are taken into account, the evidence does not support the strong lobular specialization hypothesis. Instead, it shows that lobular *rut* function is non-specialized and that STM makes use of all available functioning Kenyon cells.

## DISCUSSION

Previous studies using significance testing concluded that differences between mushroom body lobes exist that reflect functional specializations in the various memory phases (STM, MTM and LTM). These conclusions about lobular specialization included the idea that γ lobe *rut* function is sufficient for STM formation. The aim of the present study was to specifically examine the strong lobular specialization STM hypothesis. Surprisingly, the synthetic evidence is incompatible with lobular specialization, and supports the alternative idea that STM function is generalized across lobes.

Meta-analysis of strong *rut* hypomorphic alleles confirmed that they cause a 60% reduction in STM. As previously reported in the literature, the other 40% must be mediated by other molecular factors either in the Kenyon cells or elsewhere. Restoring *rut* activity with lobe-targeting drivers revealed that partial rescue occurs in both the γ and αβ lobes (mean 26% and 12%), with a partial rescue even in the α’β’ lobes (mean 6%). To rescue the majority of lost function, *rut* had to be expressed in both αβ and γ lobes (Figure 2B). These data are incompatible with the hypothesis that the lobes’ *rut* activity in the γ lobe is absolutely or strongly specialized for STM. With the synthesized evidence failing to support strong lobular specialization of *rut* in STM (Figure 2B), we considered an alternative hypothesis: that cell extent is the main predictor of a transgenic driver’s STM impact. Indeed, multivariate meta-regression models incorporating cell count show that the dominant factor influencing STM is the number of Kenyon cells targeted by a specific driver line, for both *rut* and *shi* effects (Figure 4A, B). This result refutes the hypothesis that the mushroom lobes are specialized for aversive STM function. Rather, the linear relationships lead us to conclude that the different lobes’ cells have similar potency for STM with regard to *rut* and *shi*-dependent memory processes.

Despite the paucity of experiments for *shi* in the γ, α’β’ and All lobes categories, the available data were sufficient to allow construction of a precise model of the relationship between driver cell count and memory. If STM relied on neurotransmission from a highly inter-dependent Kenyon cell ensemble, we would anticipate that *shi*^*ts*^ inhibition of small subsets of these cells would have a large effect. Instead, the observed linear trend between driver cell count and STM impact (Figure 4B) supports a model in which *shi-* dependent memory function in the αβ and γ cells occurs autonomously in individual cells or small groups of cells. It appears that strong qualitative specialization of lobular neurotransmission emerges over the subsequent minutes and hours as later memory forms [26,28].

This investigation serves as a case study in how meta-analysis and related estimation methods can help animal behavior research specifically, and biological analysis in general. Recent commentary has focused attention on reproducibility [9,35,36] and replication [37]; both of these issues are in part connected to significance testing. An encouraging aspect that was revealed as a part of this study is that the existing published data could support precise estimation with hierarchical modeling, suggesting firm data integrity. Significance testing remains the dominant statistical methodology in neuroscience and many other life sciences [38], despite its tendency to amplify interpretive variation with arbitrary thresholds. For this and other reasons, significance testing has been controversial in the behavioral sciences for half a century [39], but alternatives have yet to be implemented and adopted by the field. Estimation is an alternative data analysis framework that places the emphasis on effect sizes and the meta-analytic perspective. Exhortations in editorials and textbooks for researchers to favor estimation statistics for data analysis [10,11,40,41] have so far had little effect: neuroscience, biology, psychology and economics remain predominantly significance testing fields. This study shows how systematic review in conjunction with several meta-analytic techniques enable the synthesis of relevant available evidence so as to address inconsistencies in a field and reveal unexpected patterns in published data. Estimation as a fundamental statistical framework is also suitable for use in primary research; modern statistical texts advise that reporting effect sizes with their confidence intervals, along with the use of graphical methods, are the rightful priorities of primary data analysis [11,12,40]. Hierarchical models can similarly be applied routinely to analyze primary data with complex experimental designs, such as behavior experiments conducted in different labs [42] or by differing protocols within a lab [43,44], replacing basic methods such as ANOVA. The methods demonstrated above represent a superior statistical framework for all phases of biological research: planning, analysis, interpretation and review.

## MATERIALS AND METHODS

### Eligibility criteria and information sources

All information was sourced with searches of PubMed. To be eligible for consideration for inclusion in the systematic review each study was required to meet the following criteria: containing olfactory STM experiments on *Drosophila melanogaster* using the classic T-maze apparatus and a single training cycle **[14]**; reporting of the relevant control and experimental data as a Performance Index (PI); detailing the relevant genotypes and the number of experimental iterations (N or sample size). In addition, as STM is thought to begin to transition to MTM shortly after training **[17]**, we defined STM as using a post-training delay of 5 minutes or less. All studies selected contained transgenic manipulations of the Kenyon cells targeted to one or more of the 3 lobes (αβ, α’β’, and γ). For the systematic review of *rut* function in the Kenyon cells, studies included use of a hypomorphic allele of the *rut* gene, transgenic drivers and *UAS-rut* expression constructs. Experiments using temporally controlled expression of *rut* were excluded to eliminate the possibility of heterogeneity associated with incomplete restoration due to variations in expression longevity or strength. For the systematic review of endocytosis-dependent neurotransmission in the Kenyon cells, studies included a *UAS-shi^ts^* transgene in combination with transgenic drivers and heat treatment. Experiments that shifted *shi*^*ts*^ flies to different temperatures between training and testing were excluded to eliminate the possibility of heterogeneity due to these manipulations; only experiments using the conventional permissive-restrictive (cool-warm) comparison were included. Following the lead of the great majority of the STM literature, we did not attempt to analyze the acquisition, storage and retrieval phases of STM.

### Database search

The systematic literature search was conducted as follows and is shown as a diagram in Figure 1A. On the 11th July 2013, the search phrase ((((Drosophila) AND (learning OR memory)) AND (mushroom OR Kenyon)) AND (“2000”[Date - Publication]: “3000”[Date - Publication]) NOT review[Publication Type] was used to query PubMed, and the resulting 279 records were downloaded as two .nbib files. These files were imported into Papers2 software, and then exported as EndNote .xml. This file was loaded into EndNote X4, copied into Excel, and then imported into Apple Numbers with all bibliographic information including Title and Abstract stored in one row per record. This was then used to screen the records’ titles, abstracts and was also used to record the results of the full text screen and the detailed experimental design screen.

### Study selection

We designed the literature selection process to identify experiments that examined aversive olfactory STM (testing five minutes or less after training) in *Drosophila* as observed in the classic T-maze apparatus. We further aimed to focus the analysis on the two kinds of experiments most commonly used to understand the role of the three mushroom body lobes and the mushroom body intrinsic neurons (Kenyon cells). The first type of experiments was the usage of transgenic *rutabaga* (*rut*) to restore adenylyl cyclase function to one or more lobes in *rut* mutant flies; the second type included experiments targeting transgenic temperature-sensitive SHIBIRE (SHI^TS^) protein to the lobes to disable dynamin-dependent neurotransmission. The SHI^TS^ proteins form part of the dynamin endocytosis complex and poison its function when flies are transferred to the restrictive temperature **[45]**. The exact odor pairs under investigation were explicitly disregarded in this analysis; rather, experiments containing the full variety odor pairs were included to enable us to arrive at the most general conclusion about mushroom body function.

Two investigators (TY and JMW) performed the literature review independently and discrepancies were resolved collaboratively with a third investigator (ACC). The 279 records yielded from the PubMed search were screened in four stages to systematically exclude studies: title review, abstract reading, full text scan and a detailed review of experimental design. This process is described in Figure 1A; we used title and abstract information to discover a set of *Drosophila* behavioral studies that were likely to include aversive olfactory conditioning in adult fly (n = 65 studies) and then scanned these full text articles to find *rutabaga* restoration or *shibire*^*ts*^ experiments in the MB lobes. The final stage in the selection (“Experimental Design” in Figure 1) excluded three studies that did not meet the eligibility criteria listed above: one did not use or report an isogenic permissive control **[46]**; a second did not report sample sizes and used a post-training interval of 15 minutes **[28]**, i.e. 10 minutes later than the original criterion and 12 minutes later than other studies included; a third used pharmacogenetic temporal control of *rut* restoration **[47]**.

### Data item extraction

Two investigators (TY and JMW) extracted data independently using the measuring tool in Adobe Acrobat Pro; any discrepancies between the two extractions were resolved collaboratively. The following data were collected from each of the included experiments: author, year of publication, figure and panel numbers, genotype, mean Performance Index (PI) **[48]** with corresponding SEMs and the number of experimental iterations (N) for each mean PI value for each intervention and its related control group. To calculate STM percentages we identified a non-intervention control for each experiment, using the control that was the most similar to the experimental animals. For the *rut* restorations the closest available controls ranged from otherwise isogenic *rut*^*+*^ siblings to generic wild type (e.g. Canton-S). For the *shi*^*ts*^ experiments, including the heat-effect experiments, we used the permissive temperature controls. We also extracted experimental conditions: time delay between training and testing, odor pair, temperature, voltage, current type and relative humidity. One study’s *rut* restoration data were plotted with superimposed error bars, precluding their extraction and inclusion in the review **[25]**.

### Driver line classification

Driver lines were classified by lobe expression pattern according to the original studies themselves, except for the MB247 line which was thought to drive expression in all lobes **[21]**, but is now characterized as primarily driving expression in the αβ and γ lobes **[26,32]**. In addition, while several studies used 201Y as a γ driver, there is more recent evidence that 201Y also drives in a minority of αβ cells **[32]**; we accommodated this by doing primary analysis counting 201Y as γ, but also doing a variation in which it was counted as αβ + γ.

### Summary measures

For each experiment we calculated the intervention’s effect as a percentage change relative to the control PI. All the meta-analyses were carried out for the percentage change metric as well as the raw change in PI; the results were equivalent. We chose to report data as percentage changes for easier interpretation. The histogram in Figure 1B shows that control PI scores vary considerably across experiments; using a percentage change re-scales the phenotypes to each experiment’s wild type memory. A percentage not only reports how far a phenotype is from wild type memory but also sets a lower bound (0% memory). The standard error of each percentage change was calculated using the delta method approximation **[49,50]**.

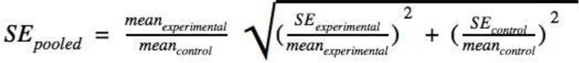

### Synthesis of results

Review Manager software {ReviewManagerRevM:wk} was used to perform nine meta-analyses: six on the *rutabaga* data, three on the *shibire* data. One random effects model meta-analysis was carried out for each mushroom body lobe and any available combinations; within each meta-analysis a subgroup analysis was performed for each driver line, except for the *rut* mutant and heat effect controls analyses, where genotype subgroups were used. Table 1 gives full details, Review Manager file is provided as Supporting Dataset 1. Complete forest plots of the six *rut* and four *shi* meta-analyses are shown in Figures 5-13. No meta-analysis was possible for *rut* restoration to the γ lobes as only one published experiment was found. Subgroup analysis of the driver lines was pre-specified. The I^2^ statistic was used as a measure of the percentage contribution of heterogeneity to the total variance in each meta-analysis, including subgroup heterogeneity **[51]**. For ease of interpretation, summary plots showed learning as a percentage of wild type learning; these were calculated by addition of the impairment effect size to 100%. We report p-values from a two-sample t-test with unequal group variances in the *rut* and *shi* summary plots, and from a t-distribution transformation for the cell count regression. Otherwise, percentage effect sizes and their 95% confidence intervals were used to interpret all results **[11]**. All 95% confidence intervals are given in the form: [95CI lower, upper].

**Figure 5.**
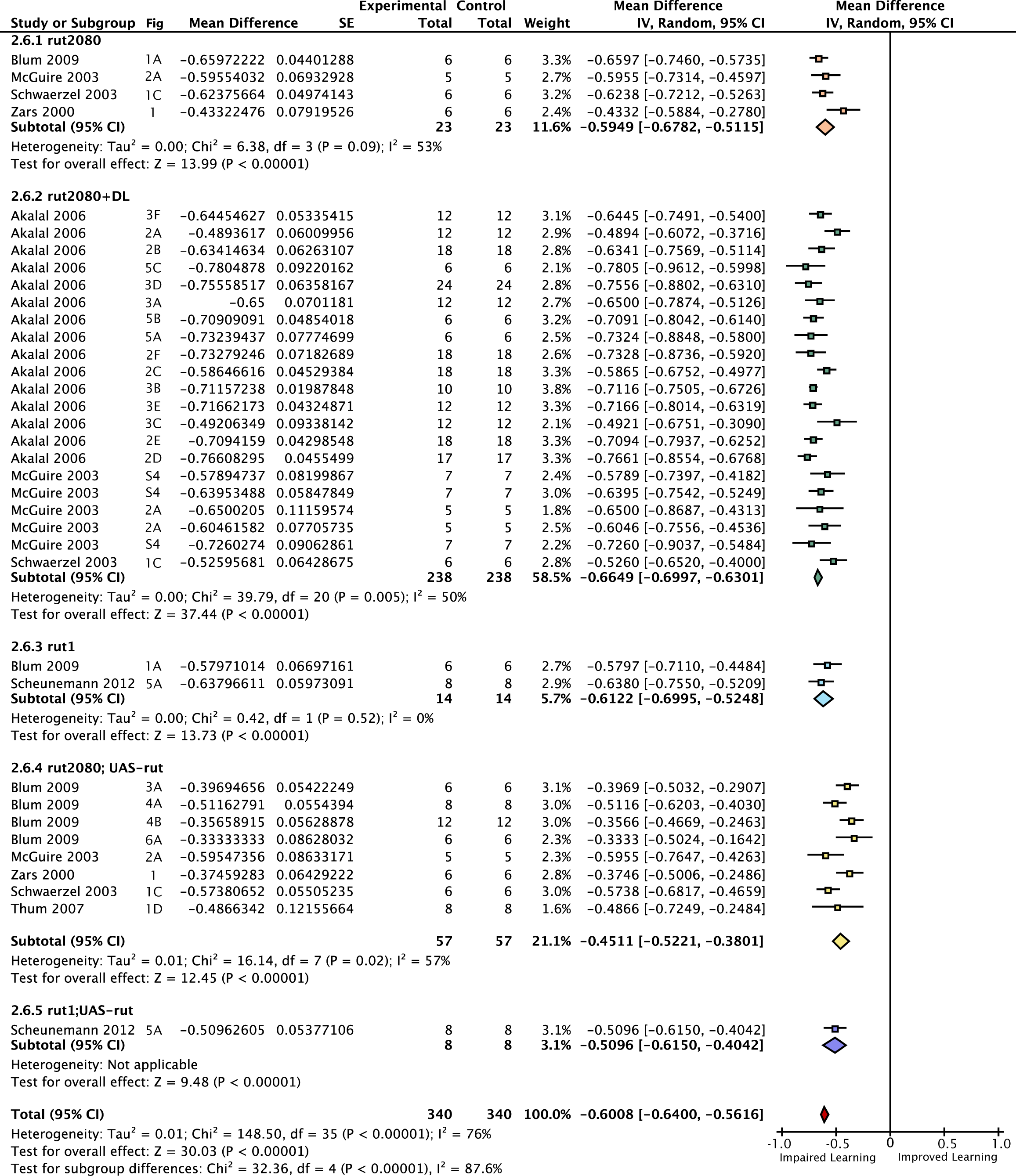
Forest plot of *rut* mutant learning changes. Each data set is identified by the source article and figure panel. This figure is a detailed version of the same plot in the main article, but uses proportional reductions instead of percentage changes. The subgroups are different driver lines, the red diamond indicates the overall estimated value range for the percentage change relative to control.

**Figure 6.**
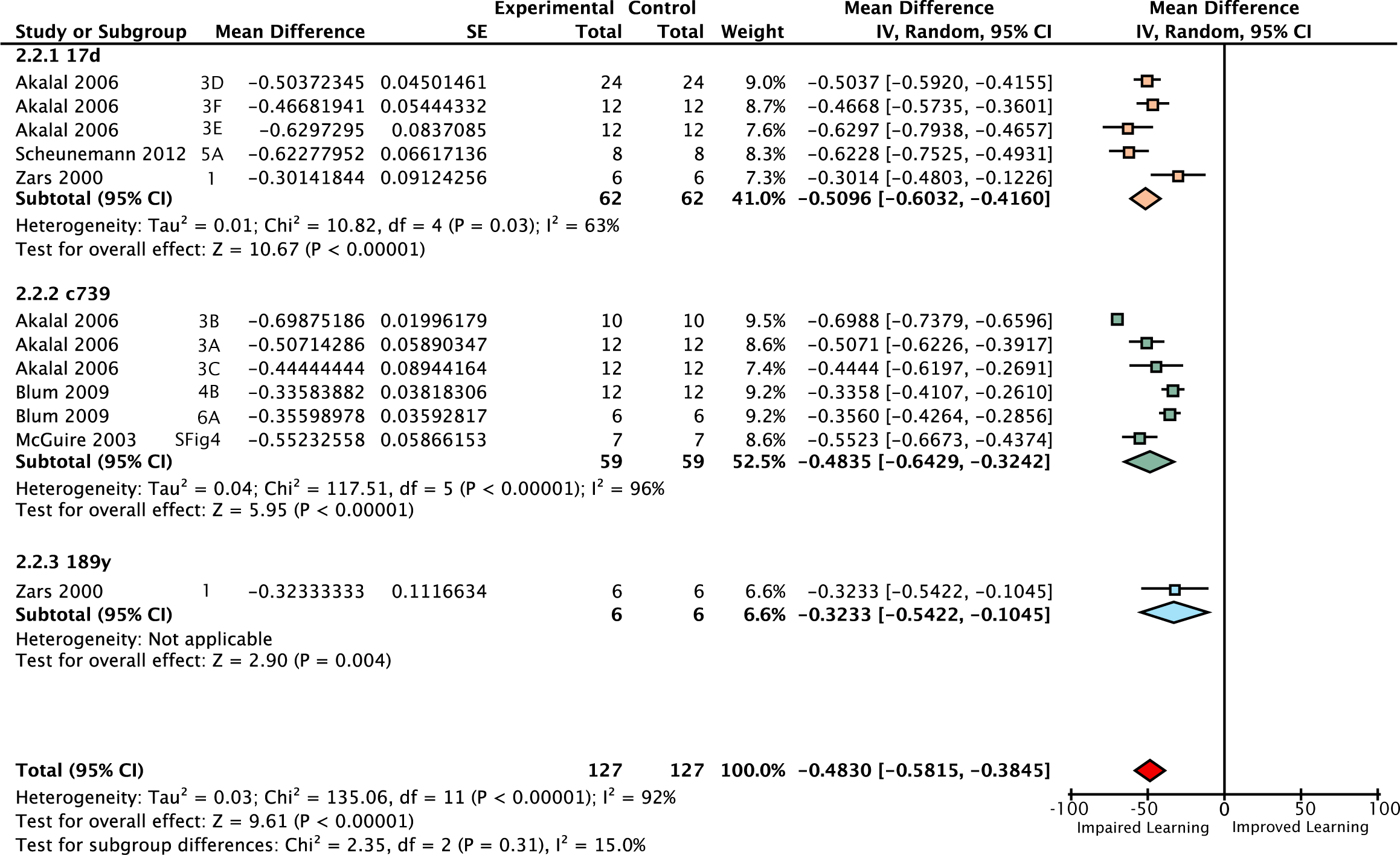
Forest plot of *rut* restoration in the αβ lobes. Each data set is identified by the source article and figure panel. The subgroups are different driver lines, the red diamond indicates the overall estimated value range for the proportional change relative to control.

**Figure 7.**
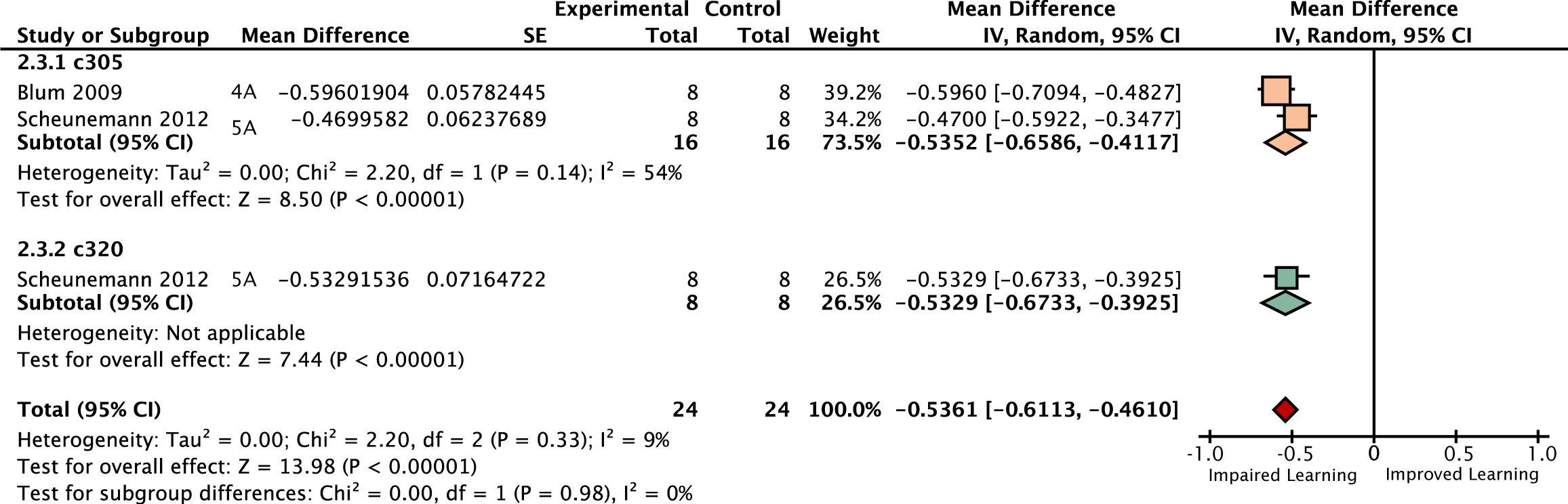
Forest plot of *rut* restoration in the α’β’ lobes. Each data set is identified by the source article and figure panel. The subgroups are different driver lines, the red diamond indicates the overall estimated value range for the proportional change relative to control.

**Figure 8.**
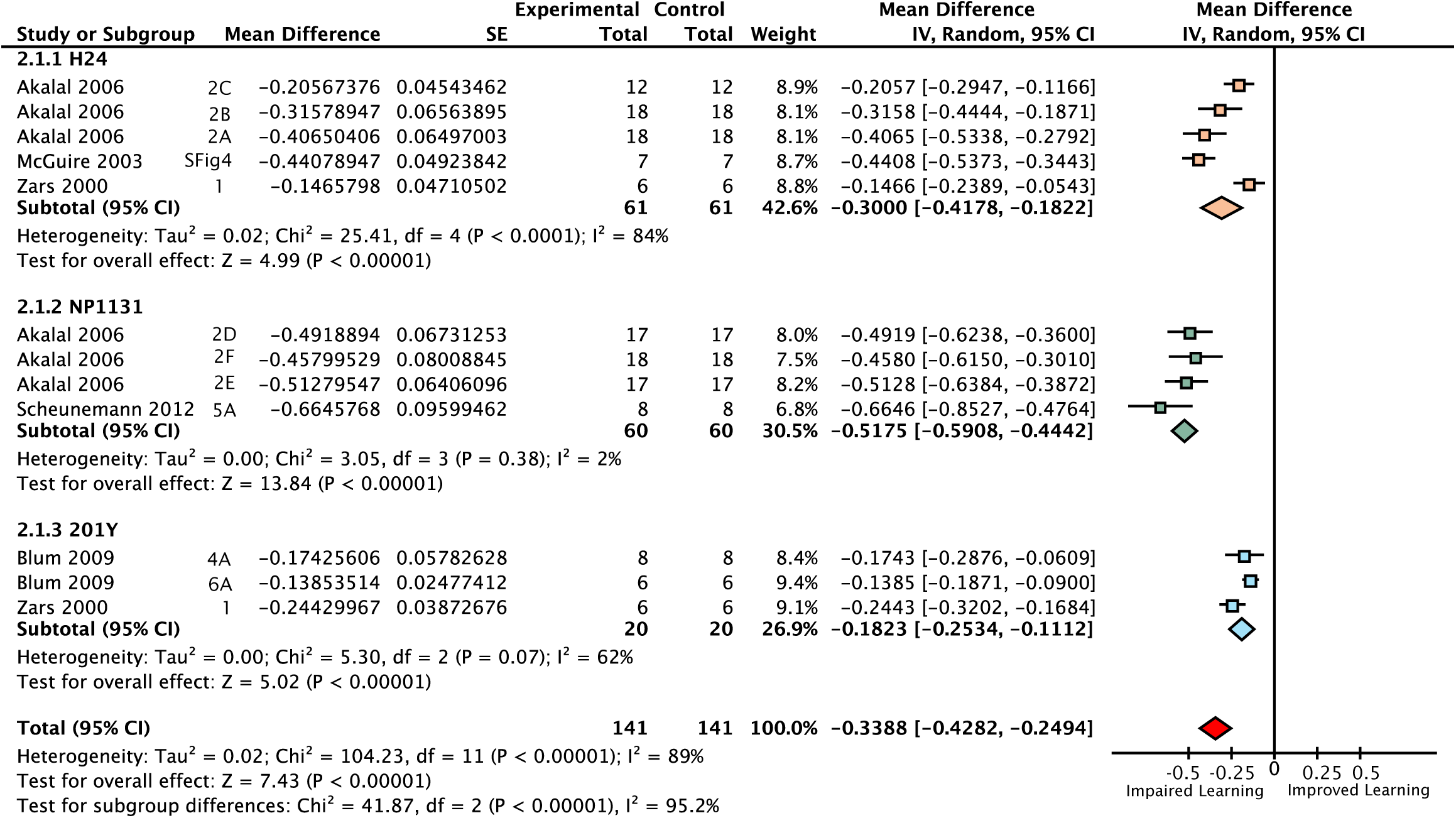
Forest plot of *rut* restoration in the γ lobes. Each data set is identified by the source article and figure panel. The subgroups are different driver lines, the red diamond indicates the overall estimated value range for the proportional change relative to control.

**Figure 9.**
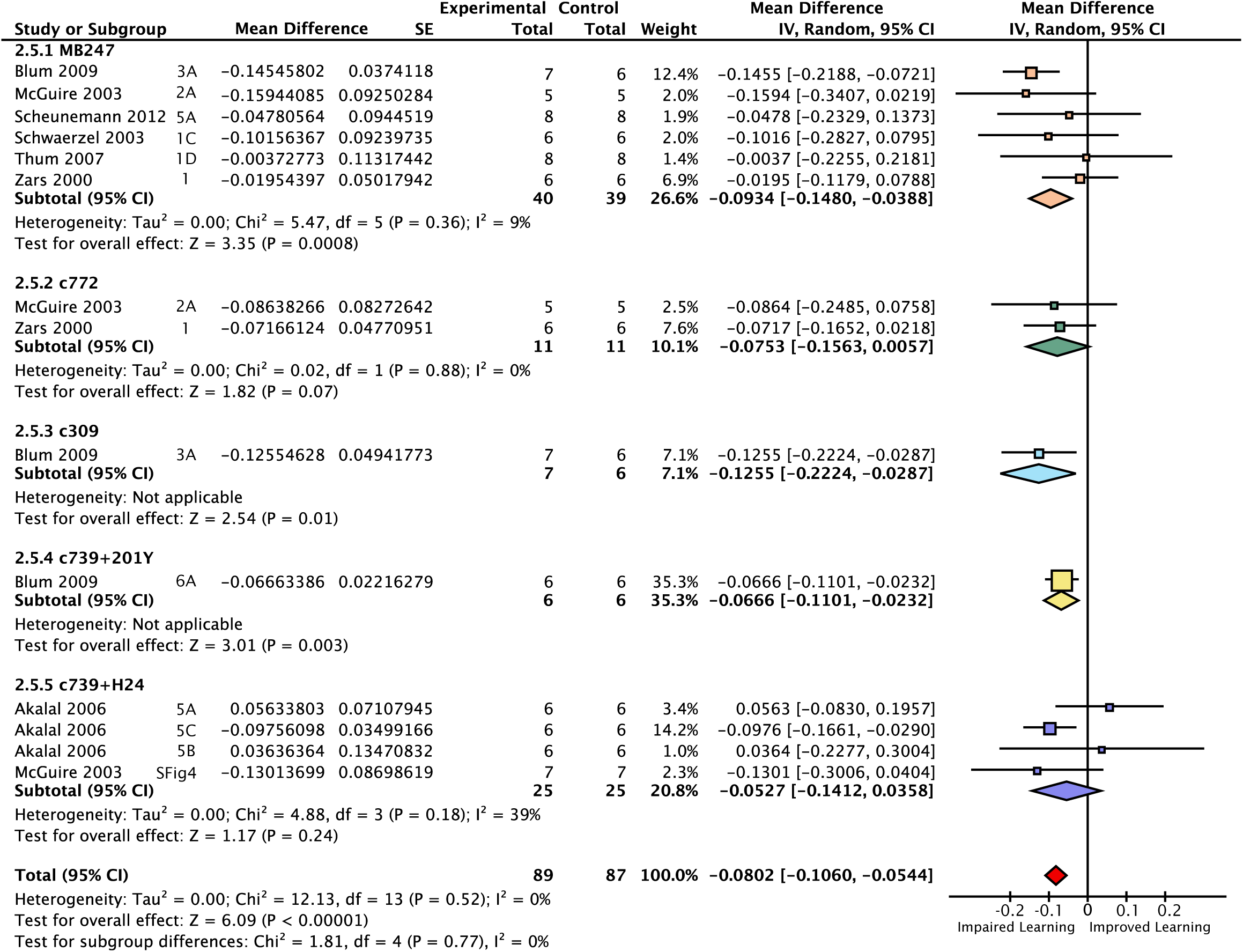
Forest plot of *rut* restoration in the αβ and γ lobes. Each data set is identified by the source article and figure panel. The subgroups are different driver lines, the red diamond indicates the overall estimated value range for the proportional change relative to control.

**Figure 10.**
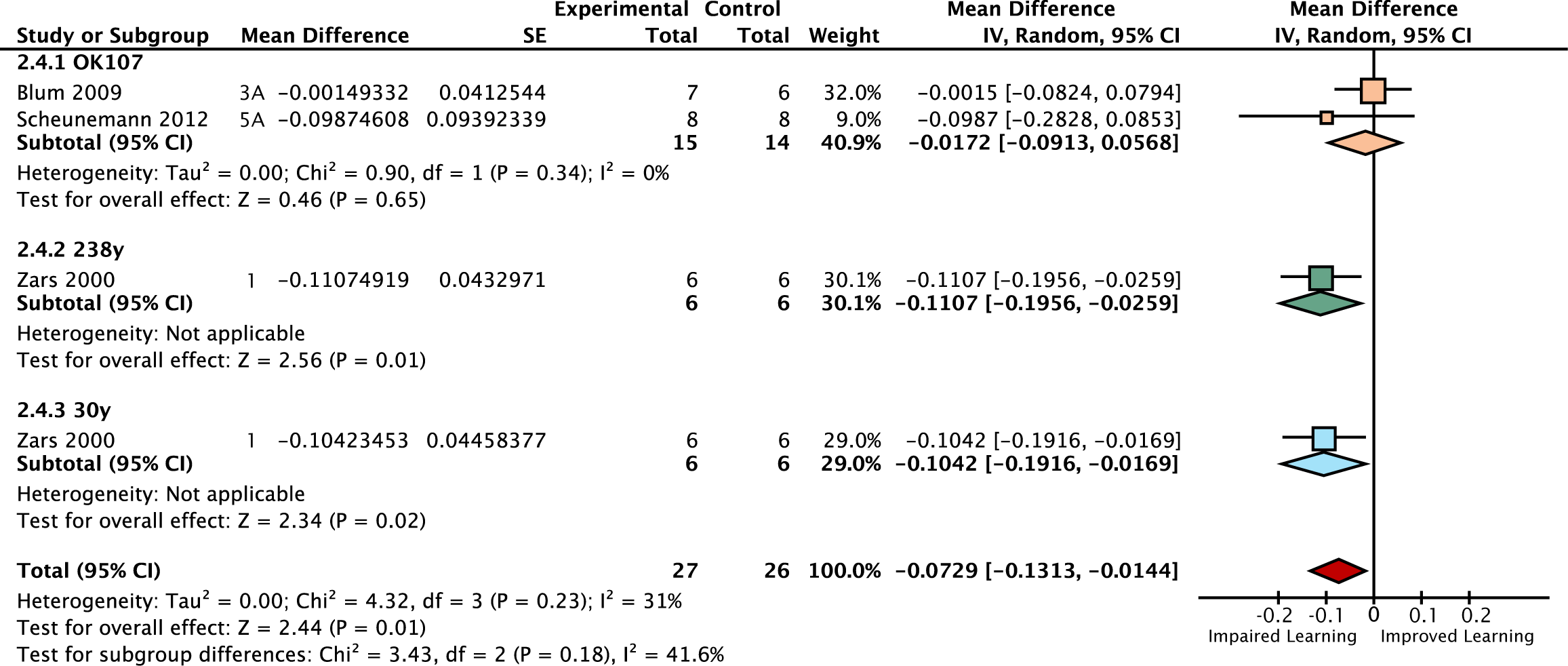
Forest plot of *rut* restoration in all lobes of the mushroom body. Each data set is identified by the source article and figure panel. The subgroups are different driver lines, the red diamond indicates the overall estimated value range for the proportional change relative to control.

**Figure 11.**
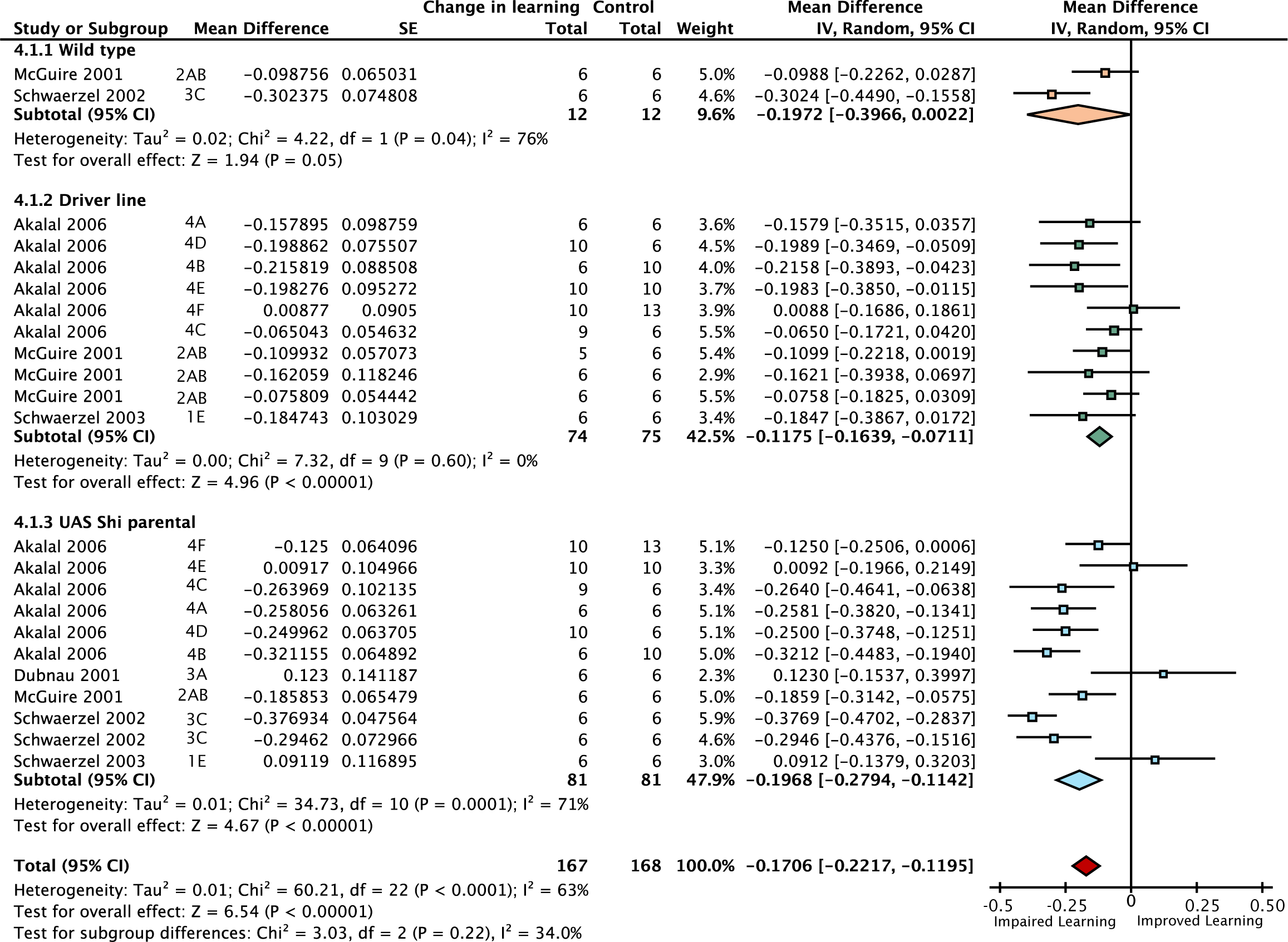
Forest plot of the effect on STM of elevating flies from permissive to restrictive temperatures. This figure is a detailed version of the same plot in the main article, but uses proportional reductions instead of percentage changes. Each data set is identified by the source article and figure panel. The subgroups are different driver lines, the red diamond indicates the overall estimated value range for the proportional change relative to control.

**Figure 12.**
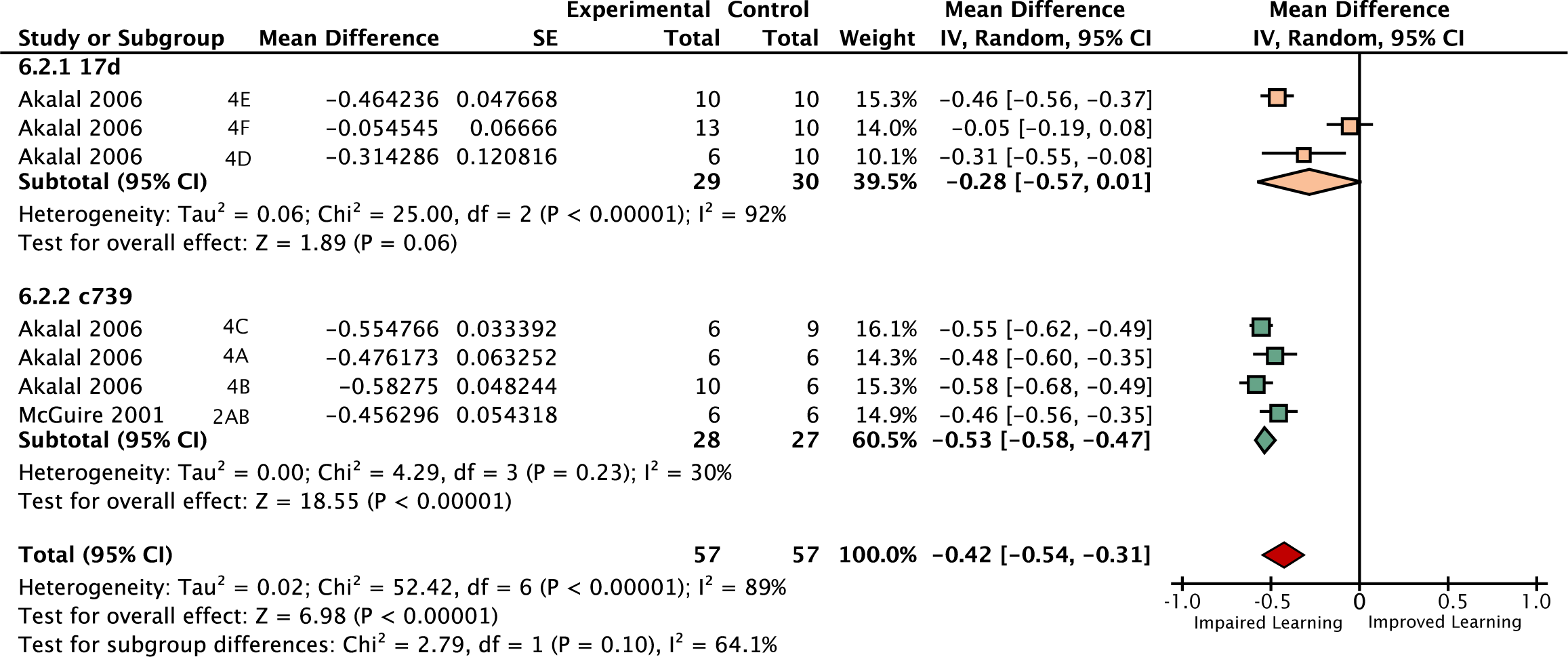
Forest plot of experiments using *shi*^*ts*^ to inactivate neurotransmission from the αβ lobes. Each data set is identified by the source article and figure panel. The subgroups are different driver lines, the red diamond indicates the overall estimated value range for the proportional change relative to control.

**Figure 13.**
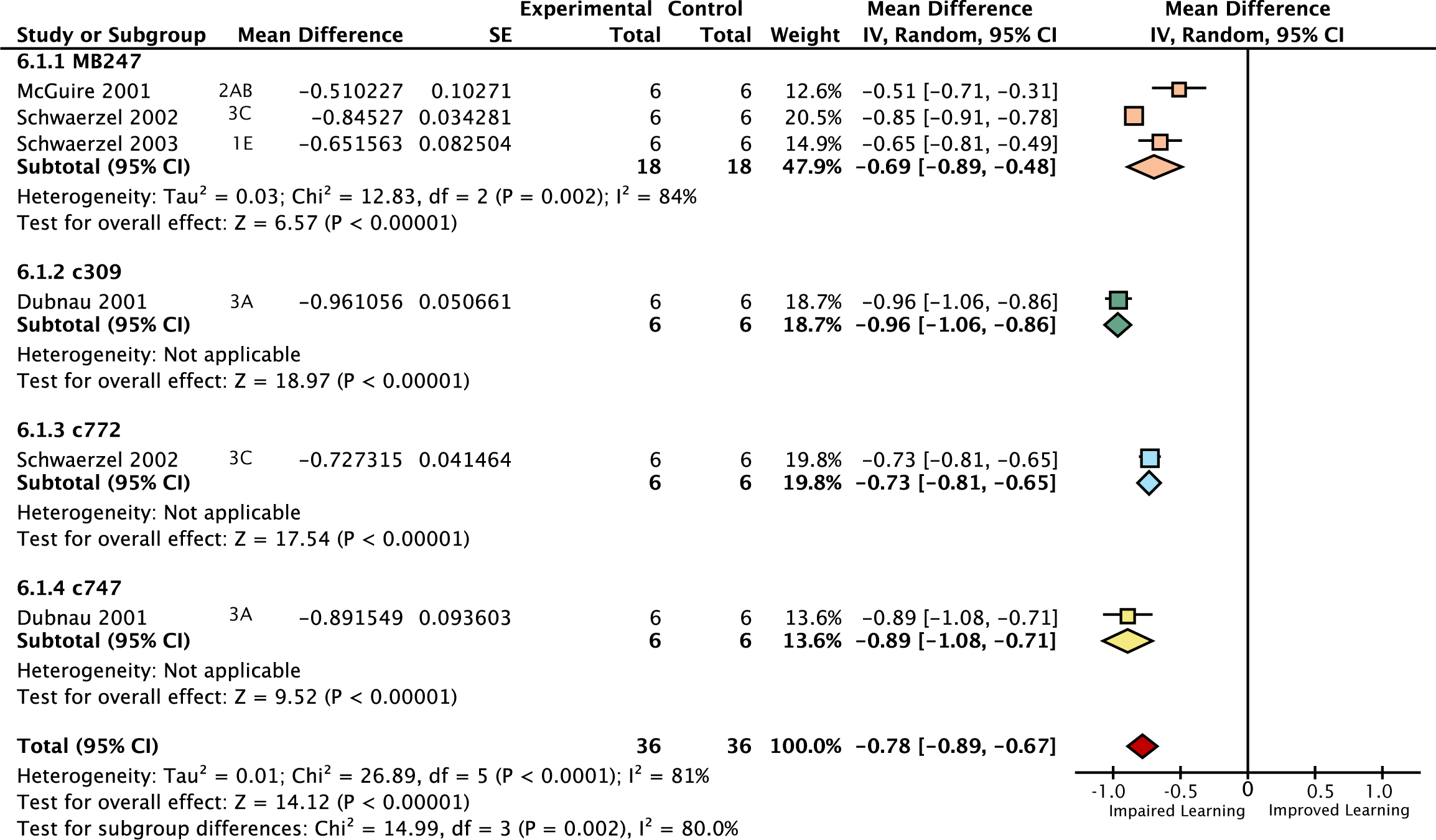
Forest plot of experiments using *shi*^*ts*^ to inactivate neurotransmission from the αβ + γ lobes. Each data set is identified by the source article and figure panel. The subgroups are different driver lines, the red diamond indicates the overall estimated value range for the proportional change relative to control.

### Meta-regression approach

Driver cell count data were extracted from a single anatomical study **[32]**. Initial examination of the relationship was done with MATLAB’s simple linear regression function (LinearModel.fit.m) on the mean values. However, this method does not account for many important aspects of the data. To accommodate the complex nature of the data, we performed multivariate hierarchical weighted meta-regression analyses of the driver effects using generalized linear mixed models (GLMM) in SAS version 9.3 software (SAS Institute, Cary, North Carolina; PROC GLIMMIX). For experiment *k* with appropriate control group *j* in study *i*, the outcome *PI*_*ijk*_ (raw change or relative percentage change) was modeled using GLMM taking into account the following:

- *The meta-analytic nature of the data:* each *PI*_*ijk*_ was estimated with a certain level of precision in the primary study/experiment. *PI*_*ijk*_were weighted in the GLMM by their corresponding precision or inverse variance (1 / *Var* (*P*_*ijk*_)) with more weight assigned to more precise *PI*_*ijk*_, as in the meta-analyses.
- *Relevant experimental design factors* (*X _ik_*)were corrected for in the GLMM to reduce the variance induced by differences in design factors between individual experiments and studies. Univariate and multivariate GLMM models were developed by including one and more-than-one design factors as independent variables in the GLMM respectively.
- *Clustering*: multiple experiments are clustered (nested) within each study and this clustering may introduce extra variability or dependence due to laboratory and personnel preferences (practice) in conducting experiments. Studies were modeled as clusters (*b*_*i*_) through a random effect with variance *τ*.
- *Shared Controls*: *rut* restorations within experiments were calculated based on a shared control, which created dependencies (correlation) between *rut* restoration effects that shared control groups. Therefore residuals (*∊*_*ijk*_) based on the same (shared) controls were correlated and residuals based on different controls were independent. Due to convergence issues arising from a paucity of data we assumed a constant correlation (*ρ*) between residuals based on the same shared controls and modeled the residual variance-covariance matrix (***∑***) with a block compound symmetry structure – blocked by shared controls, leading to conditionally independent residuals. A simple constant-variance diagonal variance-covariance matrix was used for the *shi* experiments, as matched controls were available, leading to independent residuals.

Coupling all these aspects together yielded the following univariate and multivariate weighted GLMM:

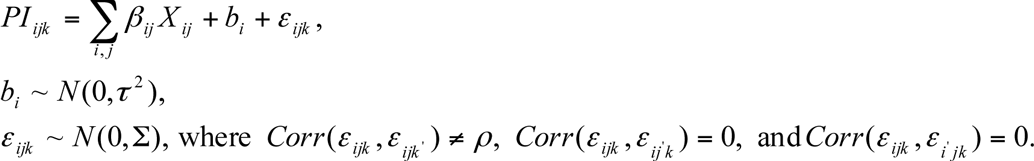

### Construction of models

Model construction started with inspection of all the available independent variables based on univariate GLMM. From Table 1, these variables included which pair of odors was used (‘ODOR PAIR’), experimental temperature (‘TEMPERATURE), delay time between testing and training (‘TIME’), shock voltage (‘VOLTAGE’), voltage type (‘AC/DC’) and relative humidity (‘RH’). The data are provided in Supporting Dataset 2. It was noted that the ODOR PAIR variable consisted of numerous categories, which would dramatically increase the degrees of freedom, so we considered replacing this with an approximation of the variable instead. Since benzaldehyde is known to stimulate gustatory receptors as well as olfactory receptors (and thus might have a different dependency on mushroom body function from other odorants), we used the presence or absence of benzaldehyde (‘BENZALDEHYDE’) as a proxy for ODOR PAIR. Of these variables, RH, AC/DC and VOLTAGE were both censored in a large proportion of experiments, and (for non-censored experiments) had mainly trivial and non-statistical effects on learning; these variables were excluded from subsequent models. TIME and BENZALDEHYDE data were available for all experiments. For *rut* experiments, both variables showed substantial and statistical influences on learning (TIME generalized-R^2^ = 0.26 [95CI 0.15, 0.36]; BENZALDEHYDE generalized-R^2^ = 0.28 [95CI 0.17, 0.39]), so these were incorporated into further multivariate meta-regression models. For the *shi* experiments, only TIME had a substantial influence on learning outcome (TIME generalized-R^2^ = 0.12 [95CI 0.04, 0.21]). Multivariate GLMM were used to account for and extract the effect of the relevant independent variables by obtaining residuals from the respective multivariate GLMM. We calculated a residual learning effect by summarizing the residuals by drivers and rescaling them by subtracting the wild type memory reference value (*shi* = 83%; *rut* = 40%). The residual learning effect was regressed against cell counts in a linear meta-regression that was weighted by sample size (the number of experiments contributing to each driver). The learning-per-cell model was built by first dividing each driver’s effect (and standard error) by its cell counts, and then fitting a multivariate GLMM with lobe categories as the main independent variable, while adjusting for other relevant experimental design factors.

## ACKNOWLEDGMENTS

We thank Jonathan Flint, Leslie Griffith, Ajay Mathuru, Joanne Yew, Gero Miesenböck, Scott Waddell, Daniel Stettler and members of the Claridge-Chang Lab for their helpful comments on earlier versions. We also wish to thank Lucy Robinson of Insight Editing London for assistance in manuscript preparation. TY, JMW and ACC were supported by a Biomedical Research Council block grant to the Neuroscience Research Partnership and the Institute of Molecular and Cell Biology. ACC received additional support from Duke-NUS Graduate Medical School, a Nuffield Department of Medicine Fellowship, a Wellcome Trust block grant to the University of Oxford and A*STAR Joint Council Office grant 1131A008. TY was supported in part by a Singapore Pre-Graduate Award from the A*STAR Graduate Academy. PNA and ESYC are supported by a National Medical Research Council block grant to the Singapore Clinical Research Institute. JMW, TY and ACC did the systematic review; JMR and TY performed the data extraction, TY performed the meta-analyses; FM performed the linear regression analysis; PNA built the meta-regression models; ACC, PNA and ESYC guided the project; ACC wrote the manuscript with contributions from the other authors.

---

**Supporting Dataset 1. Meta-analyses of *rut* and *shi* STM experiments Review Manager file.**

The file used to calculate and plot the meta-analyses shown in Figures 2, 3, 5-13.

**Supporting Dataset 2. Spreadsheet used for meta-regression of *rut* and *shi* STM experiments.**

The Excel file used as input to SAS for model construction.

